# High standing diversity masks extreme genetic erosion in a declining snake

**DOI:** 10.1101/2023.09.19.557540

**Authors:** Andrea Vittorio Pozzi, John Benjamin Owens, Bálint Üveges, Tom Major, Ellie Morris, Stuart Graham, Anatoli Togridou, Alexander S.T. Papadopulos, Wolfgang Wüster, Axel Barlow

## Abstract

Average heterozygosity is frequently used as a proxy for genetic health, and to compare genetic diversity between species and populations. However, this measurement could be misleading if the distribution of heterozygosity across the genome is highly skewed. We investigated this pitfall in methodology using whole-genome sequencing of the adder (*Vipera berus*), a species experiencing dramatic declines in the UK. We find that mean heterozygosity in adders is notably high, exceeding that of other vertebrates typically regarded as genetically diverse. Their genome-wide distribution of heterozygosity, however, approximates a negative exponential distribution, with most genome regions showing extremely low heterozygosity. Modelling approaches show that this pattern is likely to have resulted from a recent, severe bottleneck and fragmentation most likely caused by anthropogenic activity in a previously large, interconnected adder population. Our results highlight that high standing diversity may mask severe genetic erosion when declines are recent and rapid. In such situations, whole-genome sequencing may provide the best option for genetic risk assessment and targeted conservation actions.

## INTRODUCTION

The Anthropocene is characterised by an unprecedented rate of biodiversity loss (Otto, 2018; Johnson *et al*. 2017), with anthropogenic habitat destruction and range fragmentation representing some of the major drivers (Brooks *et al*. 2002; Foley *et al*. 2005). The latter can lead to the disruption of gene flow among populations, thus promoting the formation of small and isolated populations which may be pushed to the brink of extinction by low genetic diversity, inbreeding depression, and loss of genetic adaptive potential (Keller *et al*. 2002; Frankham, 2005; Fletcher *et al*. 2018). With the advent of the genomics era, average genome-wide heterozygosity has become one of the most widely used parameters to assess the genetic diversity and genetic health of populations (Schmidt *et al*. 2021). This approach has been applied to a wide variety of taxa characterised by different conservation statuses (Díez-del-Molino *et al*. 2018). Nonetheless, empirical evidence has shown that certain demographic events have the potential to mislead conservation assessments based on average genome-wide heterozygosity (Robinson *et al*. 2021).

The adder (*Vipera berus*) has the largest geographic range among terrestrial snakes, encompassing most of northern Eurasia. It is currently classified as Least Concern by the IUCN for this reason (Munkhbayar *et al*. 2021). However, populations in many countries are highly fragmented and severely declining (Graitson *et al*. 2018; Julian and Hodges, 2019; Podloucky *et al*. 2020; Munkhbayar *et al*. 2021; Guiller *et al*. 2022). This situation is typified in the UK, where anthropogenic activities have dramatically reduced and fragmented adder habitats, to the extent that most populations currently comprise less than 10 adults, and multiple local extinction events have been recorded (Baker *et al*. 2004; Worthington-Hill, 2016; Gardner *et al*. 2019; Ball *et al*. 2020). A historical survey at the beginning of the 20^th^ Century found that the species was extremely common throughout the UK (Leighton, 1901). Subsequently, numerous studies reported a decline in adder abundance and distribution, thought to be caused by agricultural intensification in the second half of the 20^th^ Century (Cooke and Arnold, 1982, Cooke and Scorgie, 1983, Hilton-Brown and Oldham, 1991).

Within the last few decades the area of suitable habitats for the adder in the UK has undergone a 40% contraction caused by urbanisation and the intensification of the industrial and agricultural sectors. Consequently, just 7% of historical sites remain occupied by the species (Gleed-Owen and Langham, 2012; Wilkinson and Arnell, 2013). Gardner *et al*. (2019) predicted that at the current rate of decline, the majority of England’s adder populations would be extinct within the next 10 years. For these reasons, the conservation status of the adder has been recently re-evaluated to “Vulnerable” in England and “Near Threatened” in the rest of the UK (Foster *et al*. 2021).

Small population sizes, such as those that characterise adder populations in the UK, are expected to result in low within-population genetic diversity. This in turn may lead to inbreeding depression, as observed in a small Swedish adder population (Madsen *et al*. 1996, 2020). However, recent genetic work based on microsatellites suggested that UK adder populations retain surprisingly high levels of heterozygosity, comparable to those from continental European populations (Ball *et al*. 2020). On the other hand, this study also highlighted high levels of relatedness between most individuals within their respective populations. Adders in the UK therefore provide an apparent, contradictory example of high genetic diversity alongside small population sizes. These results suggest that standard heterozygosity metrics based on traditional markers may be inadequate to assess genetic threats (Fischer *et al*. 2017). To further explore this, we analysed whole-genome sequencing data (∼8x coverage) from adders representing four UK populations. Our results highlight a situation where average heterozygosity fails to detect genetic erosion caused by recent demographic changes, which has broad implications for the effective conservation genetic assessment of declining species in the Anthropocene.

## MATERIALS AND METHODS

Further details regarding the methods can be found in the Supplementary Material.

### Sampling, laboratory procedures, and data processing

Four adders were sampled from each of four focal populations (Figure 1A). Samples consisted of either ventral scale clippings preserved in ethanol or freshly shed skins (maximum one day old), which were stored in separate paper envelopes prior to DNA extraction (Bangor University ethical approval: COESE2023WW01A). Publicly available sequence datasets from two Danish adders (NCBI accession number: SAMN14333252, SAMN14333253) were also used in population structure analyses.

**Figure 1.**
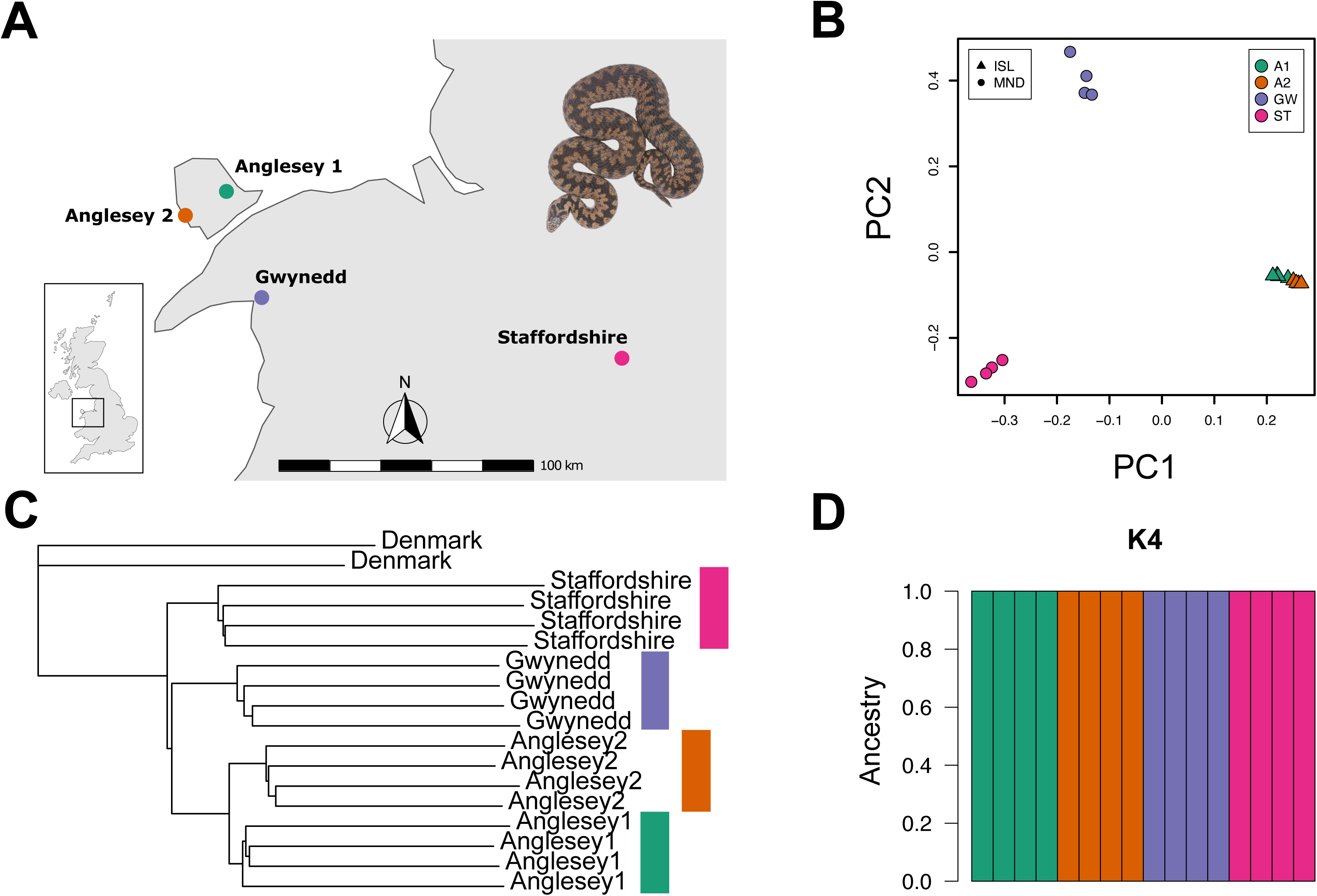
Population structure of UK adders. A. Four adders were sampled from each of the four populations from the island of Anglesey (Anglesey 1, A1; Anglesey 2, A2), mainland Wales (Gwynedd, GW), and England (Staffordshire, ST). B. Principal component analysis performed on over 7.6 million variable sites obtained from the four adder populations. Each point represents a different individual. Principal component 1 (PC1) explains 11.4 % of the total variation, while principal component 2 (PC2) explains 10.1 % of the total variation. Individuals cluster into three groups corresponding to the Staffordshire, Gwynedd and Anglesey 1and 2 populations. C. Whole genome NJ tree of the four adder populations. Genome sequences from two Danish adders were used to root the tree. The analysis reveals four distinct lineages corresponding to the four sampled populations. The four different colours represent: green for Anglesey 1, orange for Anglesey 2, purple for Gwynedd, and fuchsia for Staffordshire. D. Population structure analysis performed using NGSadmix (K=4) supports four distinct populations with no evidence of recent admixture. Each bar of the plot quantifies the ancestry proportion of each sampled genome to the respective populations. The four different colours represent the four sampled populations: green for Anglesey 1, orange for Anglesey 2, purple for Gwynedd, and fuchsia for Staffordshire.

DNA from both types of samples was extracted using a commercially available kit (DNeasy Blood & Tissue Kit, QIAGEN) following the manufacturer’s protocol. Extracted DNA was quantified using a Qubit 3 fluorometer and dsDNA HS assay kit, following the manufacturer’s instructions. Library preparation and whole-genome sequencing were carried out by an external commercial facility (Novogene UK Ltd., Cambridge), using an Illumina NovaSeq platform producing paired-end 150bp sequencing reads, aiming for 8x genome coverage.

Reads were mapped to the adder reference genome assembly (Vber.be_1.0, GenBank: GCA_000800605.1), which has a total size of 1.5 Gb, N50 value of 11.7 kb, and L50 value of 31,413. First, adapter sequences and short reads (< 30 bp) were removed using the software Cutadapt v. 1.18 (Martin, 2011). Second, overlapping pair-end reads were merged using FLASH v. 1.2.11 (Magoč and Salzberg, 2011). The mem algorithm of the software BWA v. 0.7.17 (Li and Durbin, 2009) was then used to map the processed reads to the reference genome. Finally, Samtools v. 1.3.1 (Danecek *et al*. 2021) was used to remove reads that failed to map to the reference genome, as well as secondary alignments, reads with poor mapping quality, and potential PCR duplicates. The final mapped datasets provided a mean read depth of 8.5–12.2x for UK adders, and 2.1–3.9x for Danish adders.

### Population structure

Population structure among the 16 UK adders was initially assessed through a principal component analysis (PCA). A covariance matrix was calculated in ANGSD v. 0.925 (Korneliussen *et al*. 2014) by randomly sampling a single high-quality base from the read stack of each individual at each position of the reference genome, and used to perform a PCA in R v. 4.1.2 (R Core Team, 2021). Phylogenetic relationships among the adders were investigated using neighbour-joining (NJ) analysis of a distance matrix that was calculated using the same approach in ANGSD, including the two Danish adder datasets. A NJ tree was then generated in R using the ape v. 5.6.1 package (Paradis and Schliep, 2019), specifying the Danish adders as outgroup.

Finally, genetic admixture between UK adder populations was investigated using a population clustering method based on genotype likelihoods in NGSadmix v. 32 (Skotte *et al*. 2013). Admixture proportions were calculated four times independently for three *a priori* numbers of population clusters (K=3, K=4, and K=5).

### Genetic diversity and population modelling

We investigated genetic diversity occurring at three levels: individual level, using genome wide heterozygosity; population level using nucleotide diversity (π); and between populations using Fst. Individual genome-wide heterozygosity estimation was carried out using maximum likelihood site frequency spectrum (SFS) estimations based on genotype likelihoods in ANGSD (Nielsen *et al*. 2012). We generated unfolded SFS by specifying the reference allele as the ancestral state and taking the second SFS value (positions with the ancestral and a derived allele) as our estimate of heterozygosity. We calculated the SFS along 100kb non-overlapping windows, which allowed the distribution of genome-wide heterozygosity to be examined using histogram plots in R. Individual heterozygosity values were used to calculate population averages, which were compared with published values from various vertebrate taxa (Westbury *et al*. 2018). Population-level nucleotide diversity (π), which is the average number of substitutions between two randomly sampled chromosomes in a population, was calculated along 100 kb sliding windows in ANGSD. Nucleotide diversity is zero if all sampled individuals are fixed for the same allele, representing the theoretical endpoint of population-level genetic diversity loss. Finally, we estimated weighted Fst between all population pairs in ANGSD as both a global estimate and along 100 kb sliding windows.

Population simulations were carried out to explore possible scenarios explaining the observed individual heterozygosity in UK adders using the population simulator ms (Hudson, 2002). The aim of these simulations was not to obtain precise predictions of adder population parameters and summary statistics, but rather to explore situations causing changes to the mean heterozygosity and the distribution of heterozygosity. A 50 Mb chromosome with one crossover event per generation was simulated. Estimates of genome-wide mutation rates in reptiles are not well documented, and so the human mutation rate of 1.6 x 10^-8^ mutations per site per generation was used (Lipson *et al*. 2015). Each simulation was run with 25 replicates using different starting seeds. As for the empirical data, heterozygosity was calculated along 100 kb sliding windows and histograms were used to visualise the genome-wide distribution of heterozygosity. We ran simulations involving a range of constant effective population sizes (Ne), as well as instantaneous past bottlenecks of varying severity occurring at a varying number of generations in the past.

## RESULTS

### Population structure

Ordination of individuals along PC1 and PC2 showed an overall clustering according to geographic origin, with Gwynedd, Staffordshire, and the two Anglesey populations combined, forming respective clusters (Figure 1B). The whole genome NJ tree indicates the presence of four clades corresponding to the sampling localities, differentiating the two Anglesey populations that clustered together in the PCA (Figure 1C).

Population admixture analysis grouped each individual within their respective geographic locations with no evident signal of admixture. At K=3, Anglesey 1 and Anglesey 2 clustered into a single population, compatible with the PCA result (Supplementary Figure 1A), while samples from Gwynedd and Staffordshire clustered separately. At K=4 (Figure 1D, Supplementary Figure 1B), all the samples clustered according to their geographic origin. At K=5, variance in the assignment of individuals to clusters was observed between runs, indicating insufficient signal in the data to accurately assign individuals to this number of K populations (Supplementary Figure 1C).

### Genetic diversity and population modelling

Individual mean genome-wide heterozygosity values ranged from 1.3–2.5 heterozygous sites per kb. Population averaged heterozygosity was 1.75, 1.70, 1.65, and 2.5 heterozygous sites per kb for Anglesey 1, Anglesey 2, Gwynedd and Staffordshire, respectively. These heterozygosity values are higher or comparable to those from highly diverse vertebrates (e.g. yellow baboon and chimpanzee; Figure 2A).

**Figure 2.**
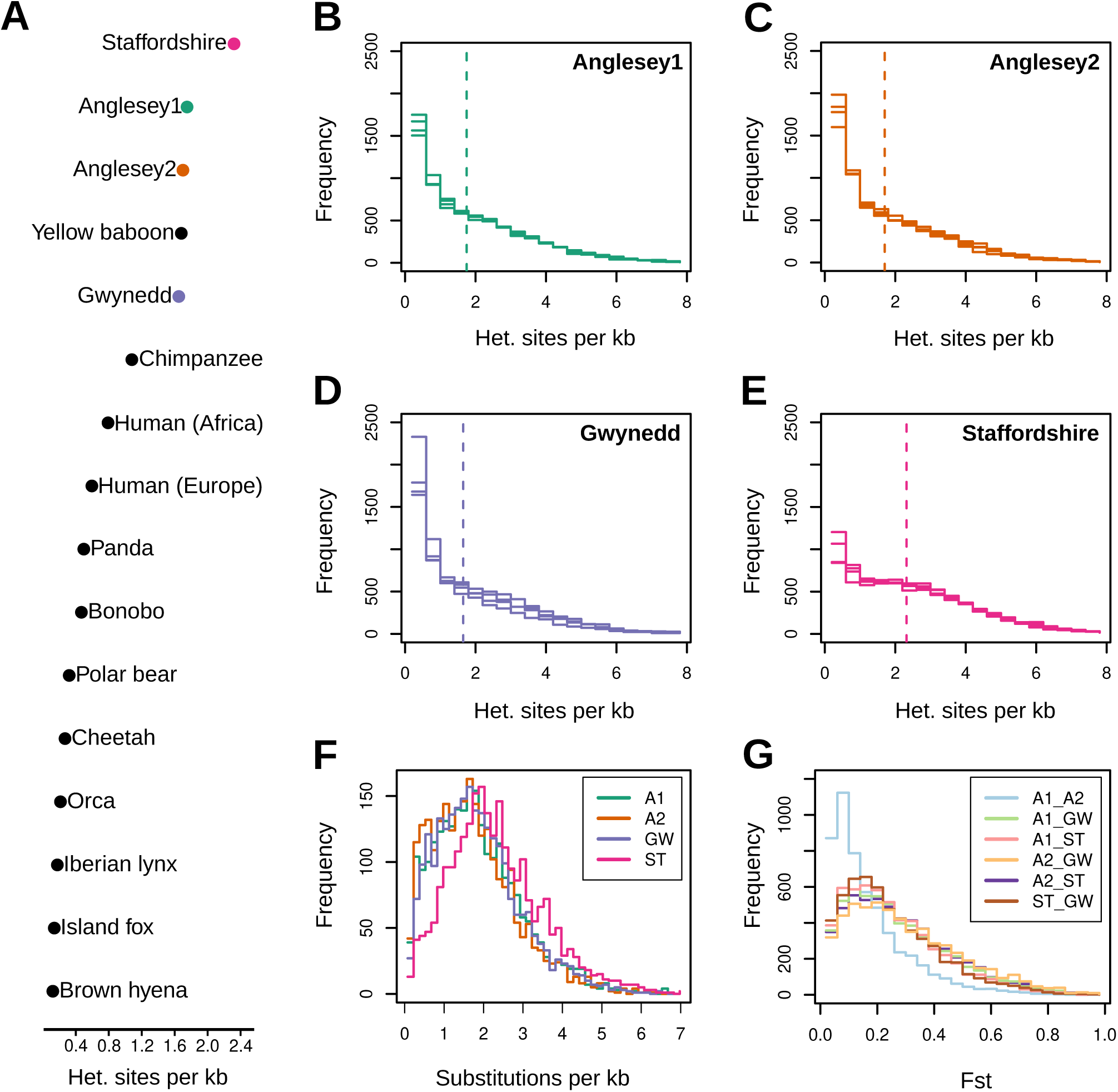
Genetic diversity in UK adders. A. Mean genome-wide heterozygosity for each population compared to values from other vertebrate taxa (obtained from Westbury *et al*. 2018), highlighting the notably high levels of heterozygosity found in UK adders. B. Genomic distribution of heterozygosity along 100 kb windows for the Anglesey 1 population. Results for the four individuals are overlaid. Dashed vertical line represent the population mean heterozygosity. C. Genomic distribution of heterozygosity along 100 kb windows for the Anglesey 2 population. Details as for B. D. Genomic distribution of heterozygosity along 100 kb windows for the Gwynedd population. Details as for B. E. Genomic distribution of heterozygosity along 100 kb windows for the Staffordshire population. Details as for B. F. Distribution of nucleotide diversity within populations along 100 kb windows. G. Distribution of Fst between-populations along 100 kb windows. Each pair of populations is represented by an overlaid line.

Examination of the distribution of heterozygosity in individual adder genomes revealed, in all cases, an asymmetric, negative-exponential distribution (Figure 2B–E). A high frequency of 100 kb windows have heterozygosity values close to zero, suggesting the presence of highly abundant runs of homozygosity (ROH) within the genomes. As a result of this asymmetrical distribution, only a small fraction of genome windows provide values equal to or greater than the mean genome-wide heterozygosity.

Analyses of population-level genetic diversity revealed differences between the studied populations. A higher proportion of genomic windows are characterised by low nucleotide diversity within the two Anglesey and the Gwynedd populations, compared to the Staffordshire population (Figure 2F). Fst analysis showed that in general a large fraction of the total genetic variation exists between populations. However, global Fst (0.16) between the Anglesey1 and Anglesey2 populations is lower than other population pairs (0.25–0.32).

Furthermore, Anglesey 1 and Anglesey 2 are characterised by a higher abundance of low-Fst windows than other populations (Figure 2G).

The population simulations (Figure 3) revealed that very large constant population sizes are required to maintain the levels of average genome-wide heterozygosity observed in the assessed UK adders (Figure 3A–D). For example, under the model’s assumptions, a Ne of 30,000 is required to achieve an average heterozygosity around 2 heterozygous positions per kb. Moreover, at these large population sizes, the distribution of heterozygosity is always approximately symmetrical and normal. Simulating a 10-fold instantaneous population bottleneck on the largest population size tested (Ne = 30,000) had little effect on mean heterozygosity or the distribution of heterozygosity within 30-60 generations post-bottleneck. More severe bottlenecks involving a 100-or 1000-fold reduction in population size, in contrast, produced an asymmetrical distribution by increasing the frequency of low diversity windows within 30 generations, similar to the distributions of heterozygosity observed in UK adders (Figure 3E–L). Notably, the most severe and recent bottleneck (1000-fold reduction, 30 generations, Figure 3E) produced a profoundly asymmetric distribution while retaining approximately 50–75 % of the original standing genetic variability (Figure 3H).

**Figure 3.**
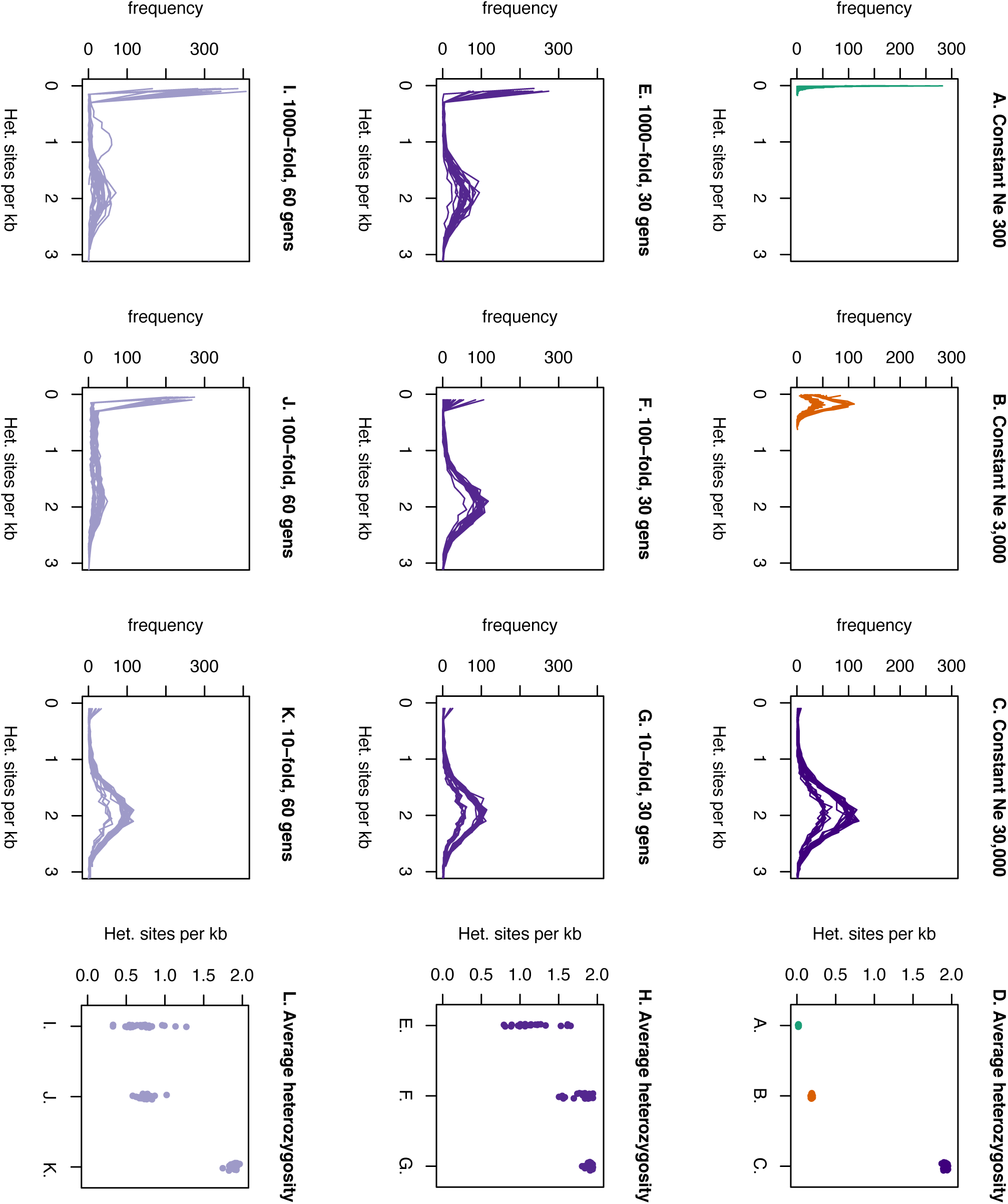
Population simulations. Population simulations support the requirement for very large population sizes in maintaining mean heterozygosity levels, and that a rapid and recent bottleneck can induce the type of asymmetric distribution of heterozygosity observed in UK adders. A. Distribution of heterozygosity along a 100 kb sliding widow for two 50 Mb chromosomes sampled from a population with a constant effective size (Ne) through time of 300. Twenty-five replicates were carried out and are represented as individual lines. B. Constant Ne of 3,000. C. Constant Ne of 30,000. D. Mean heterozygosity (y-axis) for simulations shown in A.–C. (x-axis). A random jitter has been applied to the x-axis coordinate to improve visualisation. E. Simulated bottleneck on a population with constant Ne of 30,000 (C.), involving a 1,000-fold reduction in size, sampling the population 30 generations post-bottleneck. Note the y-axis is rescaled compared to A.–C. F. 100-fold reduction in size, sampling 30 generations post-bottleneck. G. 10-fold reduction in size, sampling 30 generations post-bottleneck. H. Mean heterozygosity for simulations shown in E.–G. I. 1,000-fold reduction in size, sampling 60 generations post-bottleneck. J. 100-fold reduction in size, sampling 60 generations post-bottleneck. K. 10-fold reduction in size, sampling 60 generations post-bottleneck. L. Mean heterozygosity for simulations shown in I.–K.

## DISCUSSION

Average genome-wide heterozygosity has become one of the most widely used parameters to assess populations’ genetic diversity and genetic health (Schmidt *et al*. 2021). According to this metric, adders in the UK are highly diverse and genetically healthy, which contrasts with reports of widespread population isolation and decline (Gardner *et al*. 2019, Ball *et al*. 2020).

Through the use of whole genome sequencing, we have uncovered a pattern of profound genetic erosion in UK adders where the majority of the genome appears devoid of genetic diversity. Critically, the resulting strongly asymmetric distribution of genome-wide heterozygosity confounds genetic assessment based on mean values, which tend to be interpreted as the central position of a symmetrical (i.e. normal) distribution. This result has widespread importance for the use of mean heterozygosity in the conservation assessment of threatened species.

The overall high levels of genetic diversity occurring in UK adders indicates that their effective population size(s) must have been large in the past. However, for genetically diverse populations with stable long-term population size, genome-wide heterozygosity is expected to be normally distributed (Hohenlohe *et al*. 2021), as confirmed by our simulations and as observed in genomics studies of several species (e.g. Westbury *et al*. 2019; Robinson *et al*. 2019; Stanhope *et al*. 2023). The asymmetric distribution of heterozygosity observed in UK adders is therefore evidence against the long-term stability of their populations. Instead, our simulations show that this type of pattern can be induced by a recent, severe population bottleneck, and can precede the large-scale reduction in mean heterozygosity expected at the post bottleneck effective population size. A similar pattern of unexpectedly high mean heterozygosity at very low effective population size has been reported by a genomic study of the California condor (*Gymnogyps californianus*), which experienced an extreme population collapse to circa 22 individuals approximately four decades ago (Robinson *et al*. 2021). Given the severity of wildlife population declines associated with the Anthropocene (e.g. Dirzo *et al*. 2014; Grant *et al*. 2019; Finn *et al*. 2023), it seems likely that this general pattern could be widespread across a multitude of species and taxa.

Recently, a role for polyandry and non-random fertilisation in promoting and maintaining high levels of heterozygosity in small and isolated populations has been suggested (Madsen *et al*. 2023). While these processes may contribute to the high mean genome-wide heterozygosity recovered among our UK adder samples, they have been insufficient in maintaining the symmetrical genome-wide distribution of heterozygosity associated with large randomly-mating populations. Therefore, due to the severity of the bottleneck event experienced by UK populations, the aforementioned processes are unlikely to prevent future loss in genetic diversity.

Our results show that sole reliance on mean heterozygosity and other simple summary statistics may fail to detect advanced states of genetic erosion. The use of microsatellites or SNP panels that are ascertained to be variable in the focal population(s) are likely to exacerbate this issue, since low diversity regions are effectively excluded from the analysis (Lou *et al*. 2021). A further consideration is the temporal process of genetic diversity loss. In very severe bottlenecks, our results indicate that diversity is lost from genomes in a stepwise manner through the accumulation of runs of homozygosity, as the population tends towards a new equilibrium state of drastically reduced heterozygosity. Analysis of the distribution of heterozygosity using whole genome sequencing may therefore serve as a more sensitive method for detecting genetic erosion. Finally, the time-lag between population bottleneck and the completion of genetic diversity loss could be capitalised on by conservation strategy: if the effective population size of declining populations can be increased while mean diversity is still high, then a substantial fraction of the pre-bottleneck diversity could be retained.

One way of increasing effective population size is to increase connectivity between populations, which can be achieved through the introduction of habitat corridors, or assisted migrations, also known as “genetic rescue” (Bell *et al*. 2019). This approach has been successfully implemented to improve the genetic health of a highly inbred Swedish adder population, by the translocation of breeding males from a genetically diverse donor population (Madsen *et al*. 2020). Should this type of conservation strategy be more widely adopted, whole genome sequencing has the potential to not only identify imperilled populations suffering recent declines, but also inform the selection of donor sites to maximise conservation outcomes, demonstrating the possibility for bespoke genetic rescue (Whiteley *et al*. 2015; Khan *et al*. 2021; Bossu *et al*. 2023). Although our heterozygosity analyses reveal evidence of genetic erosion in all sampled adders, population-level diversity analyses (pairwise π, Figure 2F) provide further information to guide conservation action by quantifying the extent to which homozygous genome regions are becoming fixed in the population. Specifically, this process appears to be at a more advanced stage in the Gwynedd, Anglesey 1, and Anglesey 2 populations, in comparison to the Staffordshire population. Between population diversity analyses provides insight into the choice of potential donor sites selection for genetic rescue. Any genetic rescue attempt must consider potential diversity increase alongside the risk of outbreeding depression, and an overall consensus on where the optimal balance lies has yet to be achieved (Bell *et al*. 2019; Whiteley *et al*. 2015). Nonetheless, our Fst analysis (Figure 2G) suggests that translocation between Anglesey 1 and Anglesey 2 populations would provide a more modest diversity increase, in comparison to the other sampled population pairs. Recent progress in predicting the disruption of runs of homozygosity and reduction of genetic load resulting from population admixture (Bossu *et al*. 2023; Zhang *et al*. 2023) further highlights the potential for genomics technology in guiding genetic rescue strategy.

In conclusion, here we have shown how the persistence of high mean genome-wide heterozygosity in a declining species masks the advanced erosion of genetic diversity. This pattern is consistent with a historically large and genetically diverse meta-population that has recently undergone a severe population reduction. Our results draw attention to the potential inadequacy of mean heterozygosity and other simple summary statistics when used as the sole proxy to assess population genetic health, which may cause evidence of even severe genetic erosion to be overlooked. These findings highlight the potential of whole-genome sequencing for the rapid identification of genetic erosion and the consequent implementation of timely conservation actions.

## ACKNOWLEDGMENTS

This research did not receive any specific grant from funding agencies in the public, commercial, or not-for-profit sectors. Sequencing costs were in part covered by the Research Support Fund provided by the School of Natural Sciences, Bangor University. We acknowledge the support of the Supercomputing Wales project, which is part-funded by the European Regional Development Fund (ERDF) via the Welsh Government. B.Ü. was supported by a Research Support Grant of the Leverhulme Trust (RPG-2020-345).

## AUTHORS CONTRIBUTION

Conceptualization, A.V.P., A.S.T.P., W.W., and A.B.; Data curation, A.V.P, J.B.O, B.Ü., T.M. and A.B.; Software, A.V.P and A.B.; Investigation, A.V.P., J.B.O., E.M.; Visualization, A.V.P. and A.B.; Methodology A.V.P., A.B., Formal Analysis A.V.P., J.B.O., B.Ü., T.M., and A.B, Resources A.V.P., J.B.O., E.M., S.G., A.T., W.W., and A.B., Writing – Original Draft A.V.P., W.W., and A.B., Writing – Review & Editing A.V.P, J.B.O., B.Ü., T.M., E.M., S.G., A.T., A.S.T.P., W.W., and A.B., Project administration, A.V.P., W.W. and A.B.; Supervision W.W. and A.B.

## DATA AVAILABILITY

Raw sequencing data in fastq format will be uploaded to the European Nucleotide Archive (ENA) upon acceptance of the manuscript for publication.

## DETAILED METHODS

### Population descriptions

Samples were collected from four adder populations (Figure 1A):

- Anglesey 1: This population is in the east of Anglesey Island, within a landscape of agricultural, pastoral, and urbanised areas. The total surface area is less than 100 hectares. Lowland heathland characterises the habitat with high abundance of heather (*Calluna vulgaris*) and bushes of gorse (*Ulex europaeus*). Annual surveys of this population suggest it is currently small, likely comprising fewer than 10 adult adders.
- Anglesey 2: This population is located on the west coast of Anglesey Island. The total surface area exceeds 100 hectares. A coastal dune system characterises the habitat, dominated by marram grass (*Ammophila sp.*). Annual surveys of this population suggest it likely comprises more than 10 adult adders.
- Gwynedd: This population is located in the west of Gwynedd County, mainland North Wales. The area is bordered by agricultural land on three sides. The total surface area is less than 100 hectares. The habitat is characterised by heathland with high density of heather and gorse. Annual surveys of this population suggest it is currently small, likely comprising fewer than 10 adult adders.
- Staffordshire: This population is located within Staffordshire County in the West Midlands of England. The total surface area exceeds 100 hectares. The habitat is mainly represented by open heathland surrounded by woodland. Annual surveys of this population suggest it is large, certainly comprising more than 10 adult adders.

The straight-line distance between Anglesey 1 and Anglesey 2 is ∼20km. The Gwynedd population is situated ∼45 km from these. The island of Anglesey was last connected to mainland Wales between 5,800 and 4,600 years before the present (Roberts *et al*. 2011). Currently, the Menai Strait (250m at the narrowest point) separates Anglesey from the mainland. The Staffordshire population is located about 145 km from the Gwynedd population and 170-180 km from the two Anglesey populations.

### Data processing

Sequencing adapters and short reads (< 30 bp) were removed using the software Cutadapt v. 1.18, requiring a single base-pair overlap between adapter sequence and read and otherwise default parameters. Overlapping pair-end reads were merged using FLASH v. 1.2.11 with default parameters. The mem algorithm of the software BWA v. 0.7.17 was used to map the processed reads to the available adder reference genome assembly (Vber.be_1.0, GenBank: GCA_000800605.1). Reads that failed to map to the reference genome were then removed (-F 4), as well as secondary alignments (-F 256), reads with poor mapping quality (-q 30), and potential PCR duplicates (rmdup), using Samtools v 1.3.1.

### Principal component analysis

Population structure among the UK adders was initially assessed through a principal component analysis (PCA). A covariance matrix was obtained by randomly sampling a single read (-doIBS 1) at each position of the reference genome for each of the 16 UK adders in ANGSD v 0.925, with the following filters applied: only scaffolds > 100 kb were considered (-rf), singletons excluded by applying a minimum minor allele frequency (-minFreq) of 0.06, minimum allowed base quality score (-minQ 30), minimum read mapping quality (-minMapQ 30), and disregarding positions with missing data (-minInd 16).

### Neighbour-joining tree

Phylogenetic relationships among the adders were investigated using a distance-based neighbour-joining (NJ) method. Firstly, a distance matrix was obtained by randomly sampling reads (-doIBS 1) from each of the 16 UK adders and the two Danish adders in ANGSD using the following filters: only scaffolds > 100 kb were considered (-rf), singletons excluded by applying a minimum minor allele frequency (-minFreq) of 0.05, minimum allowed base quality score (-minQ 30), minimum read mapping quality (-minMapQ 30), and disregarding positions with missing data (-minInd 18). A NJ tree was generated in R using the Ape v. 5.6.1 library, specifying the Danish adders as outgroup.

### NGSadmix analysis

Genetic admixture between populations was investigated using a method based on genotype likelihoods in NGSadmix v. 32. Initially, a Beagle format file containing genotype likelihood data from all 16 UK adders was generated using ANGSD (-doGlf 2) with the following filters: SAMtools genotype likelihood model (-GL 1), allele frequency estimation (-doMaf 3), SNP acceptance threshold of 1e-6 (-SNP_pval 1e-6), only scaffolds > 100 kb were considered (-rf), minimum read mapping quality (-minMapQ 30), minimum allowed base quality score (-minQ 30), disregarding positions with missing data (-minInd 16), minimum individual sequencing depth (-setMinDepthInd 5), maximum individual sequencing depth (-setMaxDepthInd 20), and minimum allele frequency (-minMaf 0.1).

Admixture proportions were calculated in NGSadmix for three different number of clusters: K=3, K=4, and K=5. The analyses were repeated four independent times per each value of K changing the seed between each run (-seed).

### Heterozygosity estimation

Genome-wide heterozygosity estimation was carried out using maximum likelihood site frequency spectrum (SFS) estimations in ANGSD, using the reference genome sequence (obtained from the Swedish adder reference genome) to identify the ancestral allele (-anc). Estimations were based on scaffolds > 100 kb, with the following filters applied: minimum allowed base quality score (-minQ 30), minimum mapQ quality (-minMapQ 30), minimum individual sequencing depth (-setMinDepthInd 5), maximum individual sequencing depth (-setMaxDepthInd 24). The SAMtools genotype likelihood model was used (-GL 1). The resulting files were used to obtain the maximum likelihood estimate of SFS for 100kb non-overlapping windows.

### Nucleotide diversity within populations

In order to assess the extent of allele fixation within populations, nucleotide diversity (π) was estimated for genomic windows of 100 kb. SFS estimation for all four individuals within each population was obtained using the doSaf function (-doSaf 1) in ANGSD, refining the analysis at scaffolds > 100 kb (-rf) and using SAMtools genotype likelihood model (-GL 1). Per-site nucleotide diversity (pairwise-theta, “Pt” in ANGSD) was calculated using the saf2theta function in ANGSD, and results collected along 100kb windows (-win 100000) using the ThetaStat do_stat function.

### Fst

Estimation of the between-population weighted Fixation Index (Fst) for 100 kb windows was carried out in ANGSD. Site allele frequencies (-doSaf 1) for each population were calculated using scaffolds > 100 kb (-rf) and SAMtools genotype likelihood model (-GL 1). Then, a 2d site frequency spectrum was obtained for each pair of populations. These were used to obtain the weighted global Fst estimates (fst stats) for each population pair, as well as estimates along 100kb windows (-win 100000) (fst stats2).

### Population modelling

We carried out population simulations to explore possible scenarios explaining the observed patterns of heterozygosity in UK adders, using the population simulator ms. The aim of these simulations was not to obtain precise predictions of adder population parameters and summary statistics, but rather to explore situations explaining: 1. The occurrence of high average genome wide heterozygosity coupled with, 2. an asymmetrical distribution of heterozygosity. We simulated a 50 Mb chromosome with one crossover event per generation. Estimates of genome wide mutation rates in reptiles are not well documented, and so we used a human mutation rate of 1.6 x 10^-8^ mutations per site per generation. Each simulation was run with 25 replicates using different starting seeds. As for the empirical data, we calculated heterozygosity along 100 kb sliding windows and used histograms to visualise the genome wide distribution of heterozygosity (Figure 3).

We first explored the effective population size (Ne) required to retain the average heterozygosities observed in UK adders when the population size remains constant through time. These analyses indicated that very large population sizes are required to maintain these observed levels of heterozygosity. For example, under the assumptions applied in our model, an Ne of 30,000 is required to achieve an average heterozygosity around 2 heterozygous positions per kb, as observed in UK adders. Moreover, at these large population sizes, the distribution of heterozygosity is always approximately symmetrical and normal.

We next explored the effect of a population bottleneck on the largest population size tested (Ne 30,000). We modelled an instantaneous bottleneck of varying severities (10-, 100-, and 1000-fold reduction) at varying times in the past (30 and 60 generations in the past). These results show that, under the assumptions applied in our model, a 10-fold reduction in Ne has little effect on the distribution of heterozygosity at the timescales investigated (Figure 3 E-L). More severe bottlenecks, however, did induce an asymmetrical distribution by increasing the frequency of low diversity windows, reflecting an accumulation of runs of homozygosity along the chromosome. In general, this effect is more pronounced with increasing bottleneck severity and more distant timescales.

Overall, the population simulations demonstrate the requirement for historically large population size(s) of UK adders to explain the high observed mean heterozygosity. It is further possible to modify the genome wide pattern of heterozygosity to an asymmetrical distribution by a severe bottleneck occurring at modest timescales: i.e. within 30 generations, which is compatible with a post industrial revolution bottleneck of adders.

**Supplementary Figure 1.**
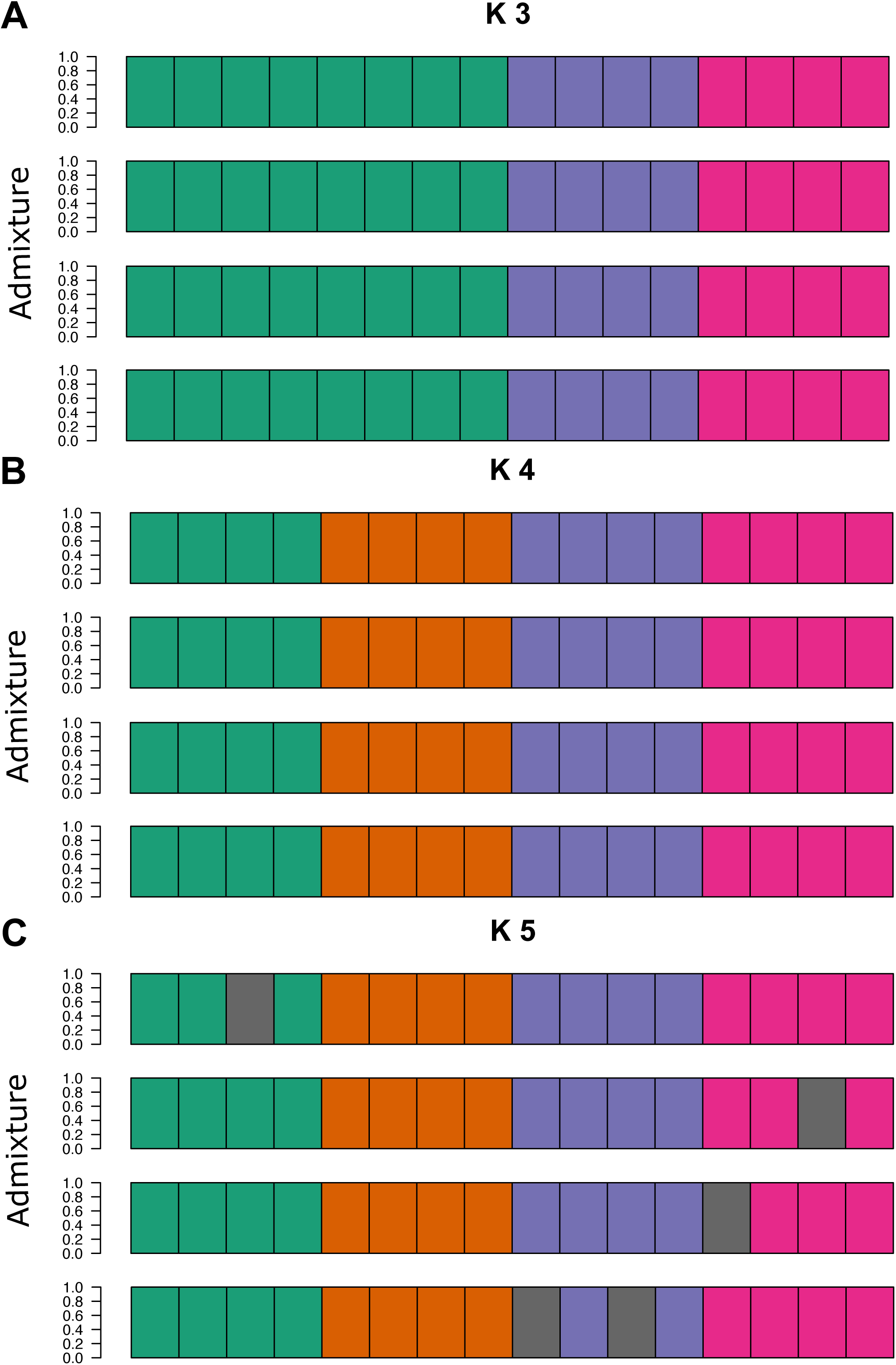
Population structure. A. Population structure analysis performed using NGSadmix and K=3. Each bar of the plot quantifies the ancestry proportion of each sampled genome to the K respective populations. Four replicated runs were performed using different seed values, and are displayed above one another. In all runs, the two Anglesey populations were clustered together, and the Gwynedd and Staffordshire populations form separate clusters. The three different colours represent: green for Anglesey 1 and Anglesey 2, purple for Gwynedd, and fuchsia for Staffordshire. B. Population structure analysis as shown in A. but with K=4. In all runs, four distinct populations corresponding to the four sampled populations were recovered. The four different colours represent: green for Anglesey 1, orange for Anglesey 2, purple for Gwynedd, and fuchsia for Staffordshire. C. Population structure analysis as shown in A. and B. but with K=5. There is a lack of consensus among the runs in terms of population structure, with some individuals variably assigned to the fifth population (grey colour, other populations coloured following D.). Overall, this result is consistent with a lack of strong sub-structuring within any the four sampled populations.

